# Coupling of replisome movement with nucleosome dynamics can contribute to the parent-daughter information transfer

**DOI:** 10.1101/154559

**Authors:** Tripti Bameta, Dibyendu Das, Ranjith Padinhateeri

## Abstract

Positioning of nucleosomes along the genomic DNA is crucial for many cellular processes that include gene regulation and higher order packaging of chromatin. The question of how nucleosome-positioning information from a parent chromatin gets transferred to the daughter chromatin is highly intriguing. Accounting for experimentally known coupling between replisome movement and nucleosome dynamics, we propose a model that can explain the inheritance of nucleosome positioning. Simulating nucleosome dynamics during replication we argue that short pausing of the replication fork, associated with nucleosome disassembly, can be the event crucial for communicating nucleosome positioning information from parent to daughter. We show that the interplay of timescales between nucleosome disassembly (*τ_p_*) at the replication fork and nucleosome sliding behind the fork (*τ_s_*) can give rise to a rich “phase diagram” having different inherited patterns of nucleosome organization. Our model predicts that only when *τ_p_* ≥ *τ_s_* the daughter chromatin can inherit the precise nucleosome positioning of the parent.

## 1. Introduction

The Fate of a cell is controlled not just by the DNA sequence alone but also by the organization and the kinetics of proteins along the DNA. In most eukaryotes, a huge fraction of the genomic DNA (more than 80% in yeast gene regions, for example) is covered by histone proteins leading to formation of a chromatin that appears like a “string of beads” [1, 2]. Advances made in the last many years have confirmed that nucleosomes and their organization play an important role in nearly all cellular processes. For example, nucleosomes are known to cover transcription factor binding sites and restrict proteins from accessing those crucial sites along the genome and, hence, regulate gene expression [3, 4, 5, 6, 7]. There are very different nucleosome organizations in coding regions and promoters of genes indicating the importance of the high diversity in nucleosome organization [3, 8, 9, 10]. Precise nucleosome organization is also crucial for higher order packaging of DNA as the polymorphic chromatin structure depends on linker length and its distribution [11, 12].

Since the precise positioning of nucleosomes is important, the natural question is how do cells transfer this information about nucleosome positioning from one generation to another? How do daughter cells know about the nature of nucleosome positioning in the parent cells? These are intriguing questions for which we do not know precise answers. One hypothesis is that the DNA sequence determines the nucleosome positioning along the genome and, hence, the information is transferred with the DNA [8, 13]. However, various experiments have indicated that the DNA sequence alone would not determine the nucleosome positioning in the genome [9, 14]—ATP-dependent chromatin remodelling, statistical positioning and other factors play equally important role [15, 16, 17, 18, 19]. Moreover, different cell types (neuronal, muscle, epithelial cells etc) have exactly the same DNA but they have very different organization of the chromatin, very different gene expression pattern and function [2]. Another major drawback of the sequence-dictated model of self-organization of nucleosomes is that attaining an “equilibrium” (steady state) nucleosome organization may take long time [20], and hence regulation of genes prior to attaining a desired nucleosome distribution may fail. An alternative hypothesis is that nucleosome positioning needs to be inherited, somehow, during replication so that the daughter cells can appropriately regulate their gene expression in an independent manner [21].

How does the de novo nucleosome assembly happen during DNA replication? Even though there have been studies on nucleosome deposition and factors associated with the octamer assembly, very little is known about the precise nucleosome organization immediately after the replication [22, 23, 24, 25, 26, 27]. There have been interesting recent experiments probing the role of nucleosome during DNA replication. Smith and Whitehouse showed that DNA replication is coupled with nucleosome assembly behind the replication fork [24]. That is, after the replication fork moves ahead and double strand formation is complete, a nucleosome is placed immediately behind the replisome machinery. They show that this nucleosome positioning is necessary for the proper progress of replication. In another paper, Yadav and Whitehouse showed that the nucleosomes behind the replication fork also gets repositioned via chromatin remodeling machines, and such remodeling is essential for obtaining certain features associated nucleosome organization [25]. Recently, Vassuer et al also studied the maturation of nucleosome organization following genome replication [26].

There has been hardly any theoretical/computational study investigating the de novo nucleosome assembly. To the best of our knowledge only the work of Osberg et al [28] investigates some aspects of the de novo assembly. However, they do not address the question of inheritance of precise nucleosome positioning from parent chromatin to the daughter.

In this work, we investigate the nucleosome organization immediately after replication, accounting for various experimentally known facts. We present a kinetic model incorporating replisome (replication fork) movement, nucleosome disassembly ahead of the fork, and nucleosome deposition and repositioning (sliding) of nucleosomes behind the fork. We show that pausing of the fork during disassembly of nucleosomes on parental chromatin and sliding/repositioning of nucleosomes on daughter chromatin behind the fork are crucial for inheritance of nucleosome positioning from a parent to daughter chromatin. We systematically explore the parameter space in the model and point out the parameter regime where inheritance of nucleosome positioning may be observed.

## Model

Here we present a model to study the nucleosome re-organization as a result of replication. In this model we start by considering an initial (parental) chromatin—DNA bound with nucleosomes—having a specific nucleosome organization. The DNA is considered a one-dimensional lattice with each base pair marked with an index *i.* The Nucleosome is modelled as a hard-core particle sitting on the lattice, occupying a space of *k* = 150 lattice sites (see Fig.1). At *t* = 0, the replisome starts replication process from the replication origin (*i* = 0), and it moves with a bare rate *v_r_* (rate of fork movement unhindered by nucleosomes) in the forward direction. As the replisome moves forward, it may encounter a nucleosome. Given that the nucleosome is a stable complex, we assume that the replisome would pause for a time *τ_p_* before it can evict the nucleosome and proceed further [29]. In other words, 1/*τ_p_* is the eviction rate of nucleosomes at the replication fork. The replisome, as it moves, creates new double-stranded DNA (dsDNA) behind it; whenever the length of the newly synthesized dsDNA is larger than the size of a nucleosome (> 150bp), a new nucleosome can occupy that space with an intrinsic rate of *k*_on_. As the replisome moves further, the process repeats. At this point, it is important to note that, as mentioned earlier, recent experiments have demonstrated that the nucleosome deposition behind the fork happens soon after the fork movement and is crucial for efficient replication [24].

**Figure 1:**
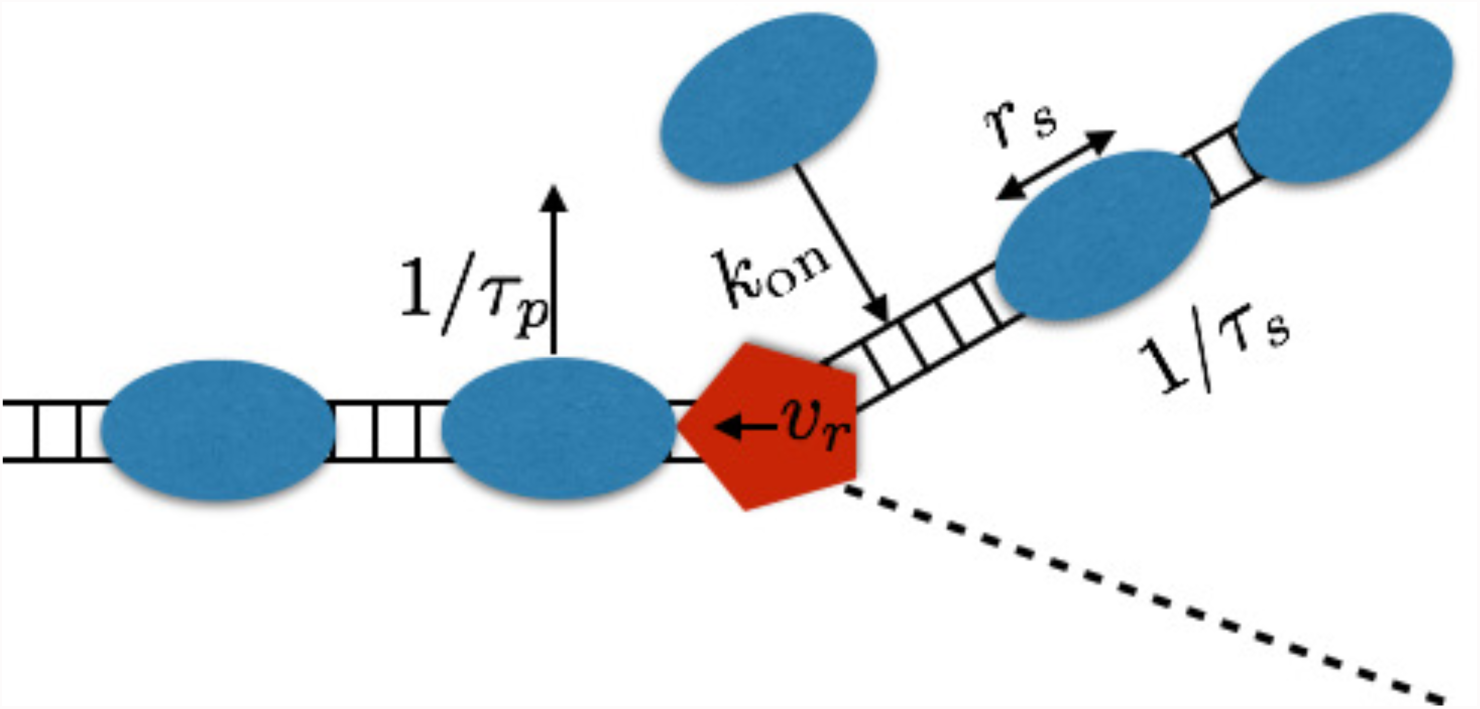
Schematic diagram representing our model for nucleosome disassembly/assembly kinetics during replication. The replisome complex (red pentagon) at the fork moves with a velocity *v_r_* in the direction shown by the arrow. As it encounters a nucleosome (blue oval) ahead, the fork movement pauses for a mean time *τ_p_* which is the timescale for the obstructing nucleosome to get disassembled (disassembly rate =1/*τ*_p_). When sufficient length of double stranded DNA (¿150 bp) is made, a nucleosome gets assembled behind the fork at a rate *k*_on_, and the newly assembled nucleosomes will get slid for a time period of *τ_s_* at an intrinsic rate of sliding *r_s_.* The sliding happens in such a way that the nucleosome will get slid to the middle of the available free region.

It has also been shown that the the newly deposited nucleosomes get slid/repositioned with the help of appropriate ATP-dependent chromatin remodelers, and this is crucial for the formation of proper nucleosome positioning [25]. In the model, taking cues from recent experiments [18, 19], we assume that a nucleosome gets slid back and forth until it settles down at the middle of the available free DNA. To achieve this repositioning, we do the following exercise: each nucleosome has a rate of sliding given by *r_s_* = *r*_*s*0_|(*i* − *i*_0_)| toward the mid position *i*_0_, from the current location *i*, with a step of size 10bp. Here *r*_*s*0_ is the intrinsic rate of sliding and *i*_0_ is the mid position of the locally available free (linker) DNA at that instant; *i*_0_ will evolve as the nearest nucleosome or the fork is displaced. However, the nucleosome does not slide for ever and stops sliding after a time *τ_s_.*

There are five rates (timescales)/parameters in the model. The rate of nucleosome fork movement, the rate of nucleosome deposition, the rate of nucleosome eviction (pause of replisome) ahead of the fork, the sliding of nucleosomes after deposition, and the duration for which the sliding will continue However, as we would describe below, many of these rates are constrained by known experimental data. For example, there are estimates for the forward movement rate of the fork [30]. In the following section we will show that the nucleosome density (occupancy) constrains the nucleosome deposition rate. Hence most of the parameters are not “free” or “independent”. In the results section we will discuss the parameters that we keep as a constant and the parameters that we systematically vary.

## Results

### A minimal model and its limitations

The simplest (or minimal) model for replication is to consider only two important processes, namely the replisome movement and the nucleosome deposition. That is, one can imagine a one-dimensional problem of a replication fork moving at a rate *v_r_* and nucleosomes being deposited behind the fork with a rate *k*_on_. This problem was considered by Osberg et al [28], and they have computed the occupancy. As a start, we also simulated replication with only these two processes, and the results are presented in the Supporting Information (SI) text. Our main findings from this simple study are (i) the average density of nucleosomes is determined by the ratio of *v_r_* to *k*_on_. (ii) Within this model, the density of the nucleosomes (the fraction of DNA covered by nucleosomes) has to be between 75% and 100%. [31, 32] (iii) The occupancy pattern in this simple model will always be uniform, one will never obtain a heterogeneous (space-dependent) nucleosome organization on an average.

The last two points mentioned above are major limitations of the minimal model. The model cannot account for even short (kilobases) regions of nucleosome density below 75%. However, we do know that there can be regions of lower density like the promoters [4, 8]. Also, in the minimal model, there is no process that gauges the space-dependence of the parental nucleosomes. To confirm this, we did a simulation where we started with a parental chromatin with a specific nucleosome positioning— the region shaded with grey in Fig.2 represents the locations where nucleosomes are positioned in the parent chromatin. Starting from this positioning, we did one iteration of replication, and the resulting average (over 500 realization) occupancy is shown in Fig.2(a)—as expected, it is a flat occupancy profile. Even though the parental chromatin had nucleosomes at specific locations, the positioning information was lost in the daughter—the reason is, within the minimal model, there is no mechanism that transfers the positional information from the parent to the daughter.

**Figure 2:**
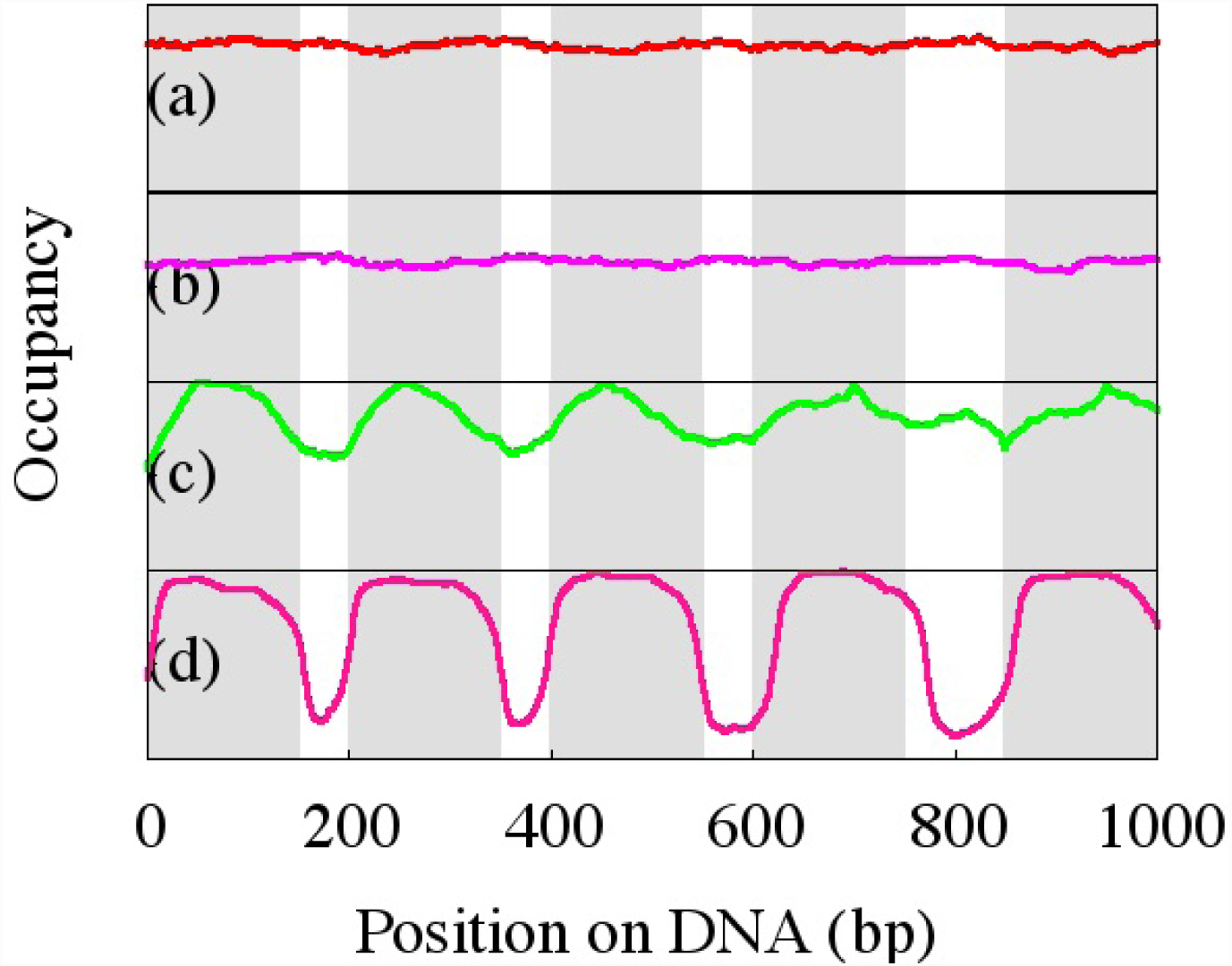
The grey region represents the fixed nucleosome positioning on the parental DNA having heterogeneous linker lengths. The resultant daughter nucleosome occupancy for various replication rules, averaged over 500 realizations, starting from the same parent: (a) Using the minimal replication model having only two parameters: *v_r_* = 500 bp *s*^−1^ and *k*_on_ = 0.1 bp^−1^*s*^−1^. (b) same as (a) but with nucleosome sliding added to the minimal replication model having sliding parameters *τ_s_* = 1*s* and *r*_*s*0_ = 1.0 bp^−1^*s*^−1^. (c) same as (b) but with fork pausing added to the minimal model having *τ_p_* = 10*s*. (d) The complete model; that is, when both pausing and sliding events are considered in addition to the fork velocity and nucleosome deposition—this replicates nucleosome positioning quite accurately. The parameter values are *v_r_* = 500 bp *s*^−1^, *k*_on_ = 0.1 bp^−1^*s*^−1^, *τ_s_* = 1*s*,*τ_p_* = 10*s*.

### Heterogeneous nucleosome organization: role of fork pausing and nucleosome sliding

In the simulations so far, we did not account for the experimentally observed [25] nucleosome repositioning (sliding). We also assumed that nucleosomes ahead of the replication fork get disassembled infinitely fast, resulting in unhindered (no pause) movement of the fork. However, in reality the replication might pause until the nucleosome ahead of the fork is removed. Given that nucleosome insertion behind the fork is strongly coupled with the movement of the fork [24, 25], we hypothesise that the timescale of such pausing, and hence the pausing in movement of the replication machinery, can be important in determining the nucleosome organization behind the fork. Therefore, we will introduce both sliding of nucleosomes behind the fork and pausing of the fork due to the removal of nucleosomes ahead of the fork. Each nucleosome, after deposition behind the fork, will be slid for a time *τ_s_* as discussed in the Model section. While the fork moves forward, as the fork reaches a nucleosome on the parent strand, the fork will pause until a time of *τ_p_* which is the time needed for clearing the way for the machinery to go forward by removing the nucleosome ahead. Since we do not know the precise values of these two parameters, we will vary them systematically and investigate the parameter regime under which one can observe experimentally sensible results. We take the bare sliding rate as *r*_*s*0_ = 1.0 bp^−1^s^−1^. The precise value of *r*_*s*0_ may not be important as long as the sliding is fast enough to take the nucleosome to its “equilibrium” position within a time of *τ_s_*.

We start our simulation with the extreme case of no pausing and consider only the sliding of nucleosomes. Then, what we have is a model with minimal moves (replisome movement+ nucleosome deposition) and the new sliding move; the results from such a model are given inFig.2(b). One can see that, with sliding and no pausing, the resulting average occupancy is homogeneous in space, and looks very different from the parental nucleosome positioning. This means that sliding alone cannot produce heterogenous nucleosome positioning. Now, we simulate the other extreme with no sliding and only pausing (see Fig.2(c)). Here, we find that the introduction of pausing brings some signature of the parental nucleosome organization. However, the occupancy pattern is not very similar to that of the parent.

Further, we simulate the model by introducing both sliding and pausing events simultaneously. First, we take the pausing timescale longer than the sliding time scale (*τ_p_* = 10s, *τ_s_* = 1s). Interestingly, in this parameter regime, the parental nucleosome occupancy is nicely replicated in the daughter chromatin strand (Fig. 2(d)). Note that even the heterogeneity in spacing is inherited in the next generation. For example, near position 200, the gap between two nucleosomes in the parent is small (≈50bp), and near position 800, the gap is large (≈100bp). One can see that in the daughter cell (even after averaging over many cells) the gap variation is nicely reproduced (Fig. 2(d)).

We have systematically studied the inheritance of nucleosome positioning by taking a few different values of *τ_p_* and *τ_s_.* In Fig.3(a), we have compared nucleosome occupancies in parent and daughter chromatins for different values of *τ_p_* and *τ_s_.* We observe that whenever both *τ_p_* and *τ_s_* are non-zero, and *τ_p_* ≥ *τ_s_* the daughter cell inherits the parent positioning reasonably well. To compare the nucleosome occupancies, we define deviation, χ, as a measure of the difference in nucleosome occupancy between the daughter and the parent,

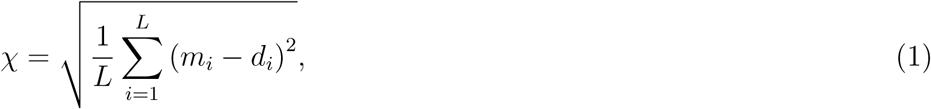

where *m_i_* and *d_i_* are occupancy of *i^th^* site in parent and daughter strand, respectively. If the nucleosome occupancy pattern between the parent and daughter is identical, then we expect the χ → 0; if the occupancy patterns are very different we expect a large value of χ close to 1. In Fig.3 (b), the deviation (χ) is plotted for different values of *τ_p_* and *τ_s_* as a heat-map with small values of χ represented by a dark violet color and large values of χ represented by a yellow color(see the colourbar on the side). This further verifies that for the parameter regime, 0 *< τ_s_ ≤ τ_p_,* the deviation is small. That is, for 0 *< τ_s_ ≤ τ_p_* the daughter faithfully inherits parental nucleosome occupancy.

**Figure 3:**
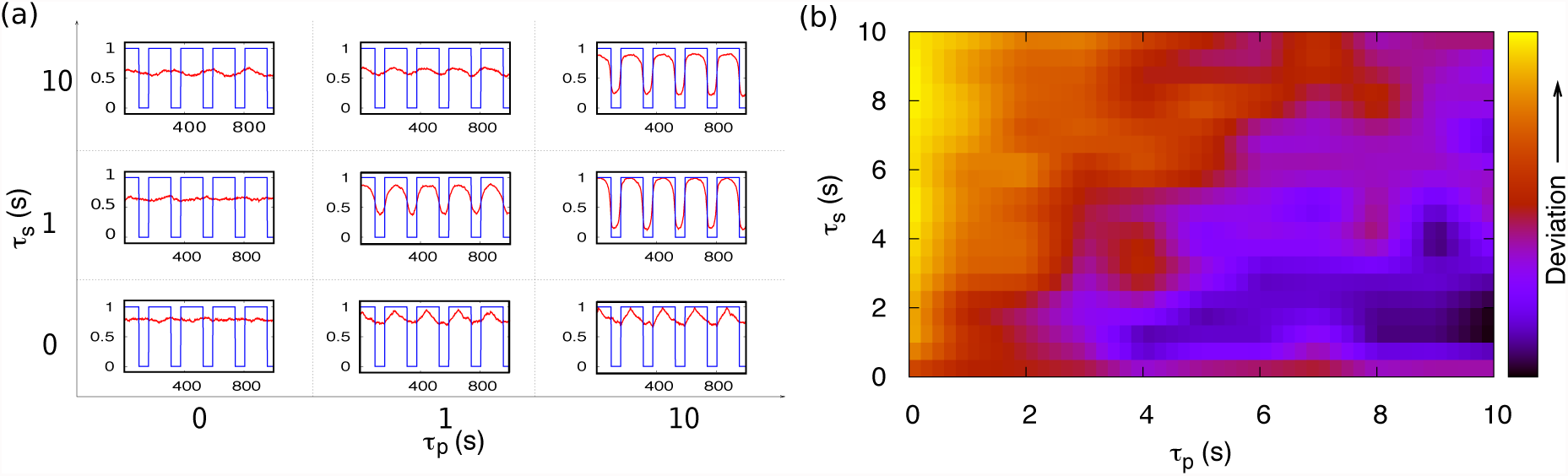
Comparison of nucleosome organization between parent and daughter chromatin for various pairs of (*τ_p_*, *τ_s_*) values. (a) Blue sharp curves represent the parental nucleosome organization and the red curves represent daughter nucleosome organization averaged over an ensemble of realizations. (b) Heat-map for the quantity ‘deviation’ (χ) as defined in eq. 1, χ increases as color varies from violet to yellow. For 0 < *τ_s_* ≤ *τ_p_* there is less deviation from parent to daughter nucleosomal organization.

### Role of strongly positioned nucleosome and barrier-like proteins

In certain parts of chromatin, it is known that there are regions where nucleosomes are “strongly” positioned, while other regions have weakly positioned nucleosomes [8, 33, 34]. Even though the DNA sequence may influence the regions with strong positioning, it is well known that factors beyond the sequence also affect nucleosome stability. For example, action (or the lack of action) of certain remodellers, histone variants (H2A.Z, H3.3), various nucleosome-binding proteins (like H1 or HMG family proteins) and histone modifications are all known to affect the stability and positioning strength of nucleosomes [35, 36, 37, 38, 39, 40]. Does stability/positioning-strength of nucleosomes have any role in transferring the nucleosome positioning information into the daughter cells?

We investigate the effect of strong vs weak nucleosome positioning and how they influence the occupancy pattern in daughter chromatin. Strongly positioned nucleosomes are defined as those nucleosomes that are more difficult to be disassembled ahead of the fork – that is, nucleosomes having a higher value of *τ_p_* are strongly positioned, while low *τ_p_* would imply weakly positioned nucleosomes. We simulate such a system with heterogeneous (high and low) *τ_p_* values 0.01s (weak) and 10s (strong) keeping *τ*_*s*_(= 1s) fixed. In a long piece of DNA, we consider two special regions with strongly positioned nucleosomes. In Fig.4(a), the two grey-shaded regions (each of length 365bp) contain two strongly positioned nucleosomes each, while the rest of the DNA has weakly positioned nucleosomes. All nucleosomes are arranged with a uniform linker length of 65bp. The resulting nucleosome positioning in the daughter cells (averaged over 5000 cells) is shown as a red curve. We observe that strongly positioned parental nucleosomes give rise to regions in daughter chromatin with high nucleosome occupancy inheriting the strong positioning. Also note that there is a positioning on either side of the strongly positioned nucleosomes implying that the strongly positioned nucleosomes can influence the positioning of the neighboring nucleosomes like in the case of the well-known statistical positioning near a strong “barrier” [15, 14, 9]. In SI text, we show that a similar inheritance of nucleosome positioning is applicable even when just one nucleosome is strongly positioned.

**Figure 4:**
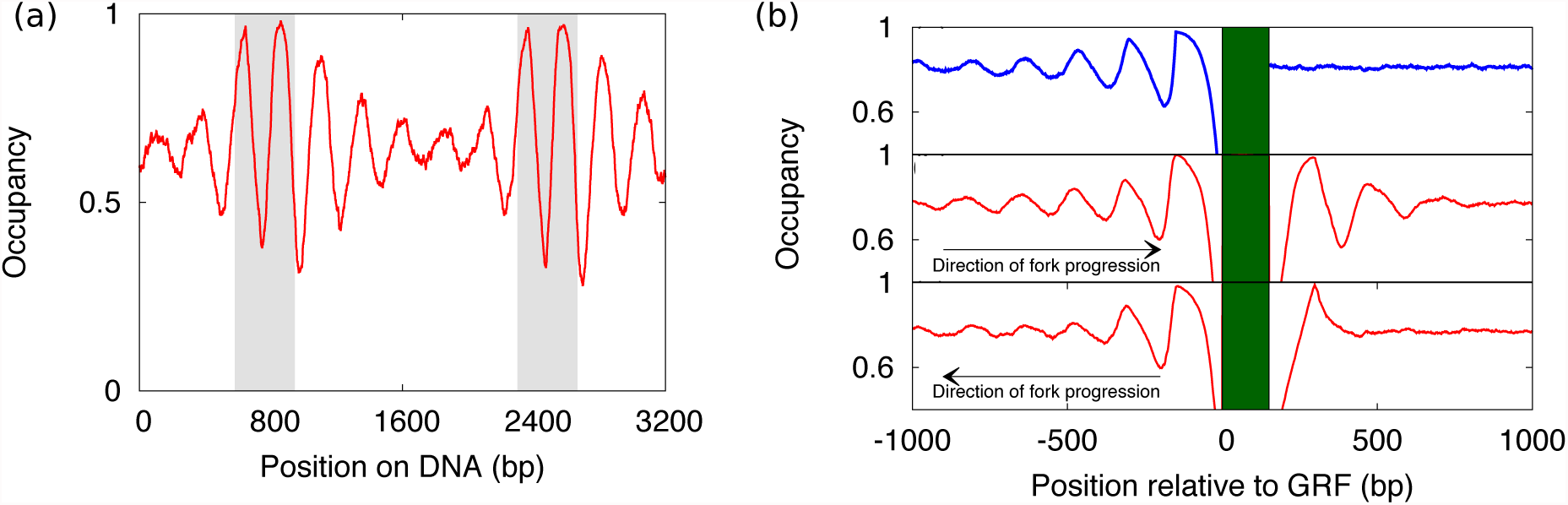
(a) The simulations are performed with parental nucleosomes in the grey shaded region (2 nucleosomes in each grey region) that are strongly positioned (*τ_p_* = 10s) and other nucleosomes that are weakly positioned (*τ_p_* = 0.01s). The daughter nucleosomal organization (occupancy) for such a heterogeneous fork pausing time is shown as the red curve. Other parameters are kept constant (*τ_s_* = 1s,*v*_*r*_ = 500 bp *s*^−1^, *k*_on_ = 0.1 bp^−1^*s*^−1^, *r*_*s*0_ = 1.0 bp^−1^*s*^−1^, Uniform linker length of 65bp.) (b) Nucleosomal positioning information transfer in the vicinity of gene regulatory factors (GRF). Top panel blue curve represents the parental nucleosome organization and the green solid bar shows presence of GRF. The middle panel shows nucleosome positioning in the daughter chromatin when the replication is performed from left to right and the bottom panel shows nucleosome positioning in the daughter chromatin when the replication is performed from right to left.

Another aspect of such local nucleosome positioning influenced by various proteins happens in the context of gene-regulatory factors(GRF). Now, we consider a situation where there is certain non-histone GRF present in the parental gene. It is known that when a bound GRF is highly stable, it can act like a “barrier” and cause statistical positioning [15, 14, 9, 41] of nucleosomes. Typically, it is known that the coding region will have the statistical positioning of nucleosomes, while the promoter regions often show different kinds of nucleosome organization [9]. How the nucleosome positioning is inherited near a GRF is an interesting question, and a recent work by Yadav and Whitehouse probed this experimentally [25]. Here we examine the prediction of our model given certain nucleosome organization reminiscent of GRF locations on the parent DNA.

On the parent DNA, on the left side of the GRF, we start with the statistical positioning of nucleosomes and on the right side with uniformly positioned nucleosomes ( flat occupancy) with mean density ≈85% (see top panel of Fig. 4(b)). We start with 5000 parent copies of the same gene, each having nucleosomes organized near the GRF in such a way that the mean of the occupancy of the parents as given in the top panel of Fig. 4(b). Each of these 5000 copy is replicated once, and we look at the nucleosome positioning on each of the gene and compute the average occupancy, which is plotted as red continuous curve in Fig. 4(b).

When we carry out the replication from left to right with regard to the GRF (in the parent, the left side has statistical positioning, the right side has uniform occupancy), we find that on the left side the statistical positioning gets replicated fairly well (see middle panel of Fig. 4(b)). However, on the right side, even though there was a flat positioning in the parent, the daughter chromatin has nucleosomes with non-uniform oscillatory occupancy in space. The physical reason for this is the following: on the left side, daughter gene inherits the parental occupancy via pausing and sliding. Whereas on the right side due to the effect of the GRF barrier, one obtains oscillatory positioning—it is well known that nucleosomes near a barrier will have spatial oscillations in occupancy. This also indicates that physical barriers will have influence near the barrier site, even with pausing and sliding. In our simulations, since the GRF is bound immediately behind the replication fork, the nucleosome depositing after the GRF “feels” (via steric exclusion) the GRF barrier and, hence, the generation of the oscillatory pattern. Please note that ATPase activity (here, sliding of nucleosomes) is an important factor in producing the oscillatory pattern as known in other contexts [9, 14].

When we carry out the replication from right to left with respect to GRF, we get the result as shown in Fig. 4(b), bottom panel. Since the machinery that is moving towards GRF is unaware of the presence of GRF until it reaches the location, the replicated chromatin will have very little influence of the barrier. However, after the GRF, the statistical positioning is reproduced. Within the short span of sliding, the nucleosome very close to the GRF feels the barrier and hence, one obtains a single peak on the right side (Fig. 4(b) bottom panel). We observe that the nucleosome organization immediately after the replication in the vicinity of GRF is tied to the replication fork progression direction (see Fig. 4(b)). This positioning may change long after replication under the influence of other events such as transcription or action of various remodellers [26]. These local remodelling events may destroy the spontaneous peak formed in Fig. 4(b) and lead to parent-like nucleosome positioning as a result of these extra events.

In an ensemble of cells, it is possible that for some cells, a particular gene gets replicated in one direction(→) while in another set of cells the same gene gets replicated in the other direction(←). This may lead to a variation in the daughter cell’s nucleosomal organization as compared to parental nucleosomal organization (see Fig. 4). Even though, in our study, we have taken a fixed barrier position, barrier location can also be varied slightly from cell to cell. This barrier can be a transcription factor or RNA polymerase itself. If we consider the barrier location as fluctuating stochastically, replication will lead to an average nucleosome positioning that may not be fully comparable to the average of the parent. Therefore, events like transcription may also play a role in “maturation” nucleosome positioning as indicated by Ref. [42].

## Discussion and conclusions

In this paper, we have addressed the question of inheritance of nucleosome organization from parent to daughter, instantly after replication, by simulating a plausible physical model. We have used various known information from published experiments and constructed a model to study the effect of different replication-related processes on nucleosome organization in daughter cells. We have first studied a bare minimum model of the fork movement and nucleosome deposition behind the fork which can only produce a homogeneous nucleosome distribution in the daughter cell irrespective of parental organization. Since the bare minimum model has no mechanism to transfer information of the heterogeneous parental nucleosome organization to the next generation, we have introduced another physically important process, which is the pausing of the replication fork on encountering a nucleosome on the parental chromatin. This interaction of the fork with nucleosomes have given some signature of parental organization in the daughter strand, but the signature has not been precise enough. Consequently, we introduced sliding of the newly deposited nucleosomes as reported in recent experiments [43, 44]. Using computer simulation we explore the parameter-space and show that when one has a finite pausing and sliding with comparable timescales, one gets replicated daughter chromatin that has similar nucleosome occupancy as that of the parental chromatin. Our model argues that strongly positioned nucleosomes act as “barriers” that will make the replication fork pause for a short period (a period comparable to the nucleosome sliding timescale) at the site of the strongly positioned nucleosomes and this pause will help transferring the positioning information from parent to daughter.

The strength of our model is that it incorporates various known experimental features such as nucleosome deposition behind the replication fork, sliding of newly deposited nucleosomes, and physically plausible events like nucleosome pausing. However, the model has various limitations or drawbacks: the first drawback is that we have not considered the extended size of replisome, which is ≈ 55bp long [45]. One reason we did not put in the size of a replisome is that, during the pause, it may happen that the replisome would partially unwrap or partially disassemble the nucleosome (which can be of a few tens of basepairs comparable to the size of the replisome) before pausing close to the dyad; this will offset the effect due to the finite size of the replisome and we will end up with a scenario that is very similar to what we have obtained here. In other words, we have not considered the size of a replisome, while we have assumed that the nucleosome at the fork will occupy full 150bp; however, the reality might be that the nucleosome may unwrap occupying only < 100bp, while the rest of the space might be occupied by the replisome. In the case of transcription, it has been reported that the RNA Polymerase pauses inside the unwrapped nucleosome near the dyad region [46]. This would be mathematically equivalent of what we did and it will not change our results. The second limitation is that the rates of processes in vivo might be very different from what we have taken for our simulations. However, we have explored the parameter-space, and found that our results will not depend on the precise value of rates; rather, the results will be true for a range of rates. The replication rate (*v*_*r*_) is constrained by known data; the nucleosome deposition rate is constrained both by the overall density of nucleosomes and the replication timescale (through the coupling). Further we have shown that any value of *τ_p_* and *τ_s_* would be fine as long as *τ_p_* ≥ *τ_s_* (Fig. 3).

Our study can be a first step in the direction of understanding the mechanism of inheritance of epigenetic information from parent to daughter, and it introduces strong physical arguments with predictive power. With our model we have been able to reproduce the parental nucleosome organization in the daughter cell with reasonable precision after disruption due to replication. While in some regions, the remodeling after replication (e.g., nucleosome rearrangement related to transcription [26]) might play some important roles, for some other regions (like heterochromatin or regions where the gene is”off”) the positioning of nucleosomes after replication may not change much. Our results will certainly be important for these latter regions. Even for regions that may change their nucleosome positioning after transcription, it is important to have a proper nucleosome positioning at all times as incorrect nucleosome positioning may expose promoters leading to unwanted transcription. We know that there are many other factors, such as DNA sequence and chemical modifications of histones, which also play significant roles in deciding the nucleosome organization. Further study is required to quantify the significance of these factors at various stages of the cell cycle.

## Acknowledgments

We acknowledge discussions with Iestyn Whitehouse. We also thank DST-INSPIRE(TB), CSIR(DD), IIT Bombay(RP) for financial support.

